# Mechanical Activation of MscL Revealed by a Locally Distributed Tension Molecular Dynamics Approach

**DOI:** 10.1101/2020.08.19.257485

**Authors:** R. R. Tatikonda, A. Anishkin, S. Sukharev, J. M. Vanegas

## Abstract

Membrane tension perceived by mechanosensitive (MS) proteins mediates cellular responses to mechanical stimuli and osmotic stresses, and it also guides multiple biological functions including cardiovascular control and development. In bacteria, MS channels function as tension-activated pores limiting excessive turgor pressure, with MscL (MS channel of large conductance) acting as an emergency release valve preventing cell lysis. Previous attempts to simulate gating transitions in MscL by either directly applying steering forces to the protein or by increasing the whole system tension were not fully successful and often disrupted the integrity of the system. We present a novel locally distributed tension molecular dynamics (LDT-MD) simulation method that allows application of forces continuously distributed among lipids surrounding the channel using a specially constructed collective variable. We report reproducible and reversible transitions of MscL to the open state with measured parameters of lateral expansion and conductivity that exactly satisfy experimental values. The LDT-MD method enables exploration of the MscL gating process with different pulling velocities and variable tension asymmetry between the inner and outer membrane leaflets. We use LDT-MD in combination with well-tempered metadynamics to reconstruct the tension-dependent free energy landscape for the opening transition in MscL.

**SIGNIFICANCE:** Membrane-embedded mechanosensitive (MS) proteins are essential for numerous biological functions including cardiovascular control and development, osmotic regulation, touch and pain sensing. In this work, we present a novel molecular dynamics simulation method that allows rapid and systematic exploration of structure, dynamics, and energetics of the mechanical transduction process in MS proteins under tightly controlled local tension distributed in the lipid rim around the protein. We provide a detailed description of the gating transition for the tension-activated bacterial mechanosensitive channel of large conductance, MscL, which is the best characterized channel of this type. MscL functions as a tension-activated emergency osmolyte release valve that limits excessive turgor pressure, prevents cell lysis and thus imparts environmental stability to most free-living bacteria.

## INTRODUCTION

Membrane-embedded mechanosensitive (MS) proteins respond to external mechanical stimuli such as pressure or tension and are involved in numerous biological functions including cardiovascular control and development (1–3), osmotic regulation (4), touch (5) and pain (6) sensing. In bacteria, MS channels such as MscS (MS channel of small conductance) and MscL (MS channel of large conductance) act as emergency release valves to prevent cell lysis under hypo-osmotic shock (7, 8). Since the initial discovery of these bacterial MS channels (9–11), numerous other membrane MS proteins have come to light including Piezo1 and Piezo2 (3, 5, 12), two-pore domain potassium (K2P) (13–15), as well as transient receptor potential (16, 17) (TRP) channels. Importantly, it is not only channels, but also G-protein coupled receptors such as angiotensin II type 1 receptors (18) (AT1R) and the newly uncovered GPR68 (2), that are apparently activated directly by tension in the lipid bilayer. It is expected that an increasing number of MS proteins will be revealed in coming years, thanks to novel high throughput screening and identification techniques (1, 2).

MscL has been extensively studied as a model system due to its high conductance, reproducible and essentially two-state behavior, small size of the pentameric complex (94.67 kDa), stability and ease of reconstitution into in-vitro systems (19–22). The availability of high-resolution crystal structures (19, 20) prompted many attempts to model (23–29) and simulate (30–41) the opening transition. However, despite the vast number of studies, the molecular mechanism of force transduction and the transition pathway in MscL and other bacterial MS channels are not fully understood. The primary stimulus for gating in MscL is membrane tension (42, 43), although externally added amphipaths, such as conically shaped lysolipids, can also induce spontaneous channel opening in the absence of net tension in the bilayer (11, 44, 45). MscL’s high activation tension-threshold (10-14 mN/m, near the membrane lytic limit) indicates a large energetic gap between the open and closed states and a substantial barrier separating the two conformations (37).

The high-resolution crystal structure of *M. tuberculosis* MscL (TbMscL) revealed a pentameric complex of identical two-transmembrane domain subunits (46). The short amphipathic N-terminal helix on the cytoplasmic side leads to transmembrane helix 1 (TM1) that lines the channel pore. The 22-residue extracellular (periplasmic) loop connects TM1 to the lipid-facing transmembrane helix 2 (TM2), which is connected through a flexible 5-residue linker with the C-terminal helix also on the cytoplasmic side. All TM1 segments tightly associate near the five-fold axis of symmetry forming a hydrophobic gate near the cytoplasmic side constricted by residues I14 and V21 in TbMscL (19–21). This hydrophobic pore is a major determinant of gating as hydrophilic amino acid substitutions in the constricted region result in channels that open under significantly lower tension or even open spontaneously (47, 48). It has been proposed that the hydrophobic pore forms a vapor-lock requiring a large tension to overcome the high surface tension of water to wet the pore and trigger a concerted opening transition (26). Disulfide cross-linking (49), FRET (50) (Förster resonance energy transfer) and EPR (37, 51) (electron paramagnetic resonance) spectroscopy studies as well as molecular sieve experiments (52) indicate a large (2.5 – 3 nm) pore opening associated with tilting of both TM helices. Beyond the pore, it is clear that gating in MscL depends on many lipid-protein and intra-protein interactions. Local stress calculations have shown that the electronegative phosphate moieties in the lipid headgroups can strongly bind positively charged residues on the N-terminal S1 domain (a highly conserved region among bacterial species (53)) that likely produces strong coupling between the membrane and protein (54). Several mutagenic studies have demonstrated the importance of this interaction as replacement or deletion of amino acids in the S1 domain leads to partial or complete loss of function (53, 55, 56). Similarly, mutations at the rim on the periplasmic side are also poorly tolerated (47, 57). This picture is further complicated by a dependence on membrane thickness (44) and lipid composition, e.g., certain anionic lipids such as phospatidyl-inositol facilitate gating in TbMscL, yet do not affect gating in the *E. coli* variant (EcMscL) (58).

Numerous theoretical and computational approaches have been put forward to understand the kinetic, structural, and thermodynamic properties of channel opening during mechanosensation. Early molecular dynamics (MD) simulations that attempted to accelerate MscL opening by exerting biasing forces directly onto the protein residues (30, 31) did not produce conformations matching the experimental data. Application of whole system tension to the bilayer in atomistic studies (32, 33) was also ineffective at inducing the large open pore predicted by experiments and typically resulted in dramatic thinning of the lipid bilayer under high tension, which questions the lateral force distribution that it transmits to the channel. Coarse-grained (CG) MD simulations with the MARTINI force-field (34–36) have succeeded in producing tension-induced gating in MscL in individual patches and pressurized liposomes, although the low-resolution model has provided limited structural information without enforcing experimentally obtained distance constraints (37, 38). Alternative approaches include finite element methods that incorporate molecular details from MD simulations into a continuum-mechanics framework to explore the mechanical transduction process between external stimulus and protein movement (39–41). At a mesoscopic level, elastic deformation models (27–29) can describe the mechanical state of the membrane-protein system and its energetics through simple terms that characterize elements such as hydrophobic mismatch and geometry, yet they lack many important atomic-level interaction details within the protein and at the membrane interface. As computational models continue to play an essential role in understanding the function of MscL and other MS proteins, there is an urgent need for a systematic approach to explore the structural, kinetic, and thermodynamic properties of MS proteins in simulation at the atomistic level.

Here, we present a novel locally distributed tension MD (LDT-MD) method that allows rapid and reversible gating of MscL in atomistic simulations by applying a local, focused at the annular lipids near the channel, but continuous tension-mimicking bias onto the membrane. Our method exploits the strong coupling at the lipid-protein interface to induce channel opening without having to directly bias protein residues (30, 31) or apply unphysically high tensions to the whole system (32, 33). LDT-MD is highly effective and achieves channel gating on the scale of a few nanoseconds without disrupting membrane structure or protein-lipid contact. We show that structural parameters such as helical tilts, pore radius, area expansion, and conductance of the LDT-MD-attained open state of MscL are in excellent agreement with experimental measurements. In combination with enhanced-sampling methods such as metadynamics, we characterize the free energy landscape of opening under tension. Furthermore, the flexibility of our method permits systematic exploration of asymmetric stimuli on the two membrane leaflets analogous to the effect of single-sided addition of lysolipids. The method is generally applicable to any membrane protein.

## METHODS

Molecular dynamics simulations were carried out using the GROMACS simulation package version 2016.3 (59). The MscL-membrane system was simulated with a combination of the GROMOS 43A1-S3 (60) force-field (FF) for lipids, GROMOS 54A7 (61) FF for proteins, and the SPC/E water model (62). Newton’s equations of motion were computed using a classical leapfrog integrator with a time step of 2 fs. Lennard-Jones interactions were calculated using a plain cutoff scheme up to a distance of 1.6 nm. Long-range electrostatic interactions were computed with the particle-mesh Ewald method using a real-space cutoff of 1.6 nm and Fourier grid spacing of 0.15 nm. Temperature was held constant at 37 °C with a Nose-Hoover thermostat and pressure was held constant at 1 atm with a semi-isotropically coupled Parrinello-Rahman barostat. The Berendsen barostat was used for constant tension simulations due to technical limitations within GROMACS. MscL from *M. tuberculosis* (PDBID 2OAR (20)) was embedded into an equilibrated POPE (1-palmitoyl-2-oleoyl-sn-glycero-3-phosphoethanolamine) membrane patch composed of 480 lipids and sufficient water molecules to maintain a layer > 1.5nm between the channel and its periodic image. The initial simulation structure (equilibrated for 500 ns) was taken from Vanegas and Arroyo (54) and further equilibrated for 200 ns. For simulations under constant tension, the membrane-protein system was equilibrated for another 150 ns with and additional number of water molecules added to maintain sufficient distance between the protein and its periodic image as the lateral area of the box increased while its thickness decreased.

### LDT simulations and free energy calculations

Conventional MD and enhanced sampling simulations with the LDT CV (Eq. 4) were conducted with a PLUMED (63, 64) (v. 2.5) patched version of GROMACS 2016. For both symmetric and asymmetric LDT-MD simulations, the contributions from lipids in each membrane leaflet were considered separately as detailed in Eq. 7. The same values of the constants *a* = 1 nm and *r*_min_ = 1.1 nm were used to compute both *ξ*_peri_ and *ξ*_cyto_. Spring constants with values of *κ* = 10,000 – 100,000 kJ · mol^−1^ · nm^2^ were used for constant velocity simulations with linearly moving harmonic restraints (Eq. 6).

Free energy profiles of MscL gating under tension (10, 25, and 35 mN/m) and without tension were obtained using well-tempered metadynamics (65) with multiple walkers (66). In this method, a time dependent bias potential of the form

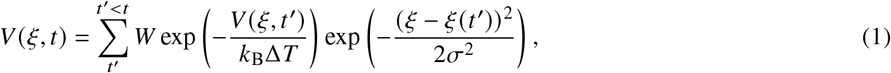

where *W* and *σ* are the height and width of the added Gaussian hills, and Δ*T* is a fictitious maximum increase in temperature that ensures convergence by limiting the extent of the free energy exploration. In the long time limit, the unbiased free energy, *F*(*ξ*), may be recovered from

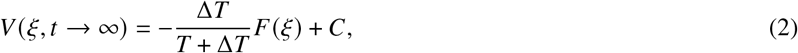

where C is an immaterial constant. The value of Δ*T* is adjusted by setting the ‘bias factor’ parameter, 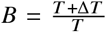 and the frequency of addition of Gaussian hills is adjusted by a fixed deposition rate, *ω*. The same values of *σ* = 0.05, *B* = 12, *W* = 1.2 kJ/mol, and *ω* = 10 ps were used for all free energy calculations. Multiple walkers (200) that simultaneously contribute to the same biasing potential were employed to accelerate sampling, where initial configurations for each walker were obtained from constant velocity LDT-MD simulations of tensioned systems followed by equilibration for 5 ns with a static harmonic potential centered at the chosen value of *ξ*. Each walker was then simulated for 30 ns during the WTMetaD run and the combined simulation time to obtain each free energy profile was 6 *μ*s. Convergence of the free energy calculations was monitored by computing the root-mean-squared deviation of the estimated profiles in 2 ns intervals (per walker).

### Data analysis

Pore radius at residue V21 was obtained from the subunit-averaged distance between the center of mass (COM) of each residue to the combined COM of all five residues in the plane of the membrane (*x* – *y*)

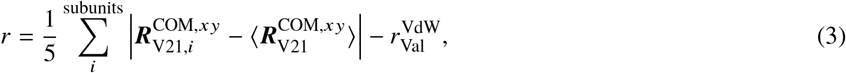

where 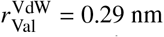 is the effective radius of a single Val residue calculated from the VdW volume of 105 nm^3^. Projected protein area was obtained by creating a 2-dimensional Voronoi tessellation of all the protein and lipid atoms according to their *x* – *y* positions. The total protein area was computed by taking the sum of the individual areas of all protein atoms. Voronoi tessellations were calculated using the `gmx_LS` voronoi utility (based on the Voro++ library (67)) part of the GROMACS-LS package (68). Vectors to compute tilt and crossing angles of the TM helices were calculated from the positions of C*_α_* of N13 and I46 for TM1 and of L69 and R98 for TM2. Conductance of the channel is estimated using the continuum approximation where the pore is considered a conductor of certain length and cross-sectional area profile taking into account the access resistance on both sides of the pore determined from the effective radii of the cytoplasmic and periplasmic pore entrances (69–71). Graphical representations of the MscL systems were created using UCSF Chimera and ChimeraX (72, 73).

## RESULTS AND DISCUSSION

### Locally Distributed Tension – rapid and reversible gating of MscL

Inspired by our earlier studies to gate MscL by directly applying forces on protein-bound lipids (54), we have developed a general LDT-MD method that mimics tension yet does not require excessive strain on the membrane. This accelerated method is based on a collective variable (CV) that weights lipids depending on their lateral proximity to the channel in the membrane plane (*x* – *y*) through a smooth hyperbolic tangent stepping function,

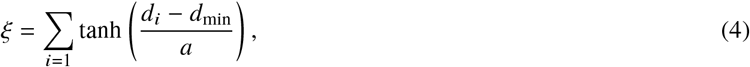

where *d_i_* = |***d**_i_*| is the lateral distance between the centers of mass of any given lipid and the protein 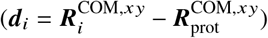, *d*_min_ is an effective minimum distance from the center of mass of the protein, and *a* is a scaling parameter that modulates how quickly the hyperbolic tangent function approaches unity (schematically shown in Fig. 1a). Similarly defined CVs have been used by others to study pore formation (74, 75), membrane bending (76), and hydrophobic peptide insertion (77).

**FIGURE 1.**
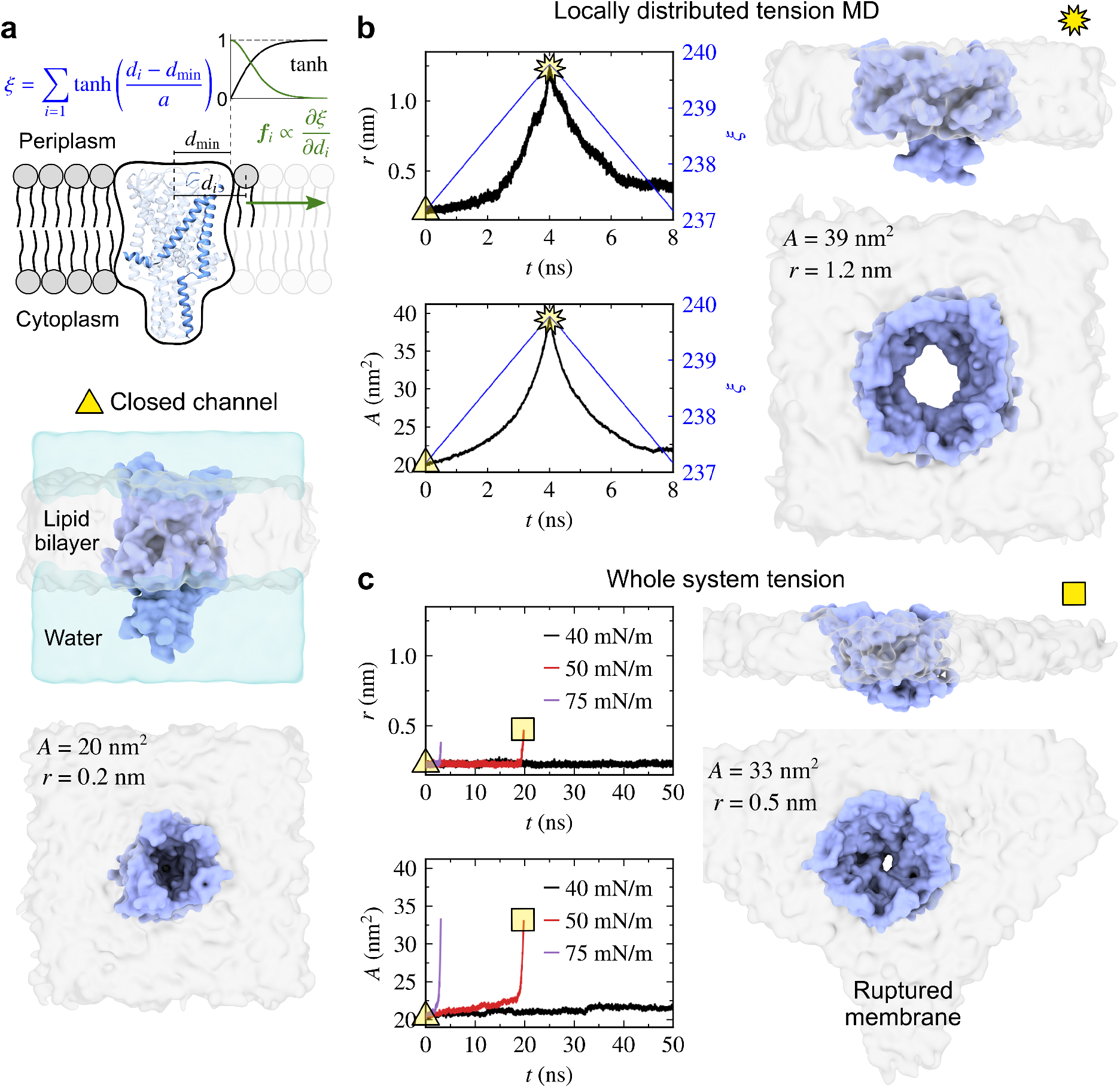
Reversible gating by tension-mimicking LDT-MD method. (a) Schematic representation of the Locally Distributed Tension (LDT) collective variable, *ξ*, based on lateral distances from MscL (see text for definition of quantities). (b) Reversible gating of MscL by linearly biasing *ξ* (blue line) with a moving harmonic potential results in changes in pore radius, *r*, (measured at residue V21, black line) and in-plane projected protein area, *A* (red line). Space filling surface representations of MscL (blue) and membrane (gray) show side and top (not showing the cytoplasmic C-terminal helical bundle to emphasize the open pore) views during various stages of gating. All surface representations are shown at the same scale. (c) Gating attempt by application of whole system, net, tensions (40 mN/m, 50 mN/m and 75 mN/m). The 40 mN/m simulation was run for an additional 100 ns without any noticeable change in either quantity. For surface tensions higher than 40 mN/m, MscL begins expanding but the destabilized membrane ruptures causing the MD simulation to crash.

Application of a CV-biasing potential, *V*(*ξ*), generates forces on all lipids,

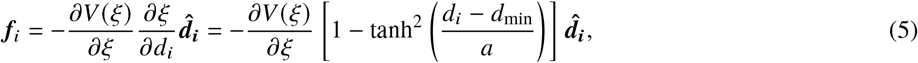

that rapidly decrease as the distance from the protein increases (Fig. 1a). We term this bias ‘Locally Distributed Tension’ (LDT) as it exerts a radially-varying distribution of lateral forces focused on the annulus of lipids that interact directly with the channel, while minimally disturbing the membrane farther away. LDT allows application of localized pulling or pushing forces, driving expansion or contraction respectively, that can rapidly gate the channel in a controlled and reversible manner as shown in Fig. 1b through the pore radius (defined by the narrowest constriction in the hydrophobic core at residue V21) and in-plane projected protein area (see Methods for precise definitions of both of these quantities). Our LDT-MD method gently drives MscL from the closed state to a large open pore (Fig. 1b) with radius ~1.2 nm in a few ns without rupturing the membrane. Reversing the biasing potential after the channel reaches an open state (*t* = 4 ns, Fig. 1b) produces an almost symmetric return to the closed configuration. If we allow the open channel to spontaneously close after LDT steering in the absence of any biasing potential, we observe a rapid contraction of the pore to an expanded state in a few hundred ps followed by a much slower return to the fully closed state over ~ 50 ns (see Fig. S1 in the Supporting Material). The absolute value of *ξ* in our simulations (Fig. 1b) is immaterial and is simply a reflection of roughly the total number of lipids in each leaflet (240).

In comparison to the LDT-MD method, tension applied to the whole system by uniformly expanding the simulation cell in the *x* – *y* plane, via the barostat, is not effective in gating MscL during timescales accessible to conventional atomistic MD simulations as shown in Fig. 1c. Application of large tensions up to 40 mN/m (3× the experimental midpoint), similarly to previous simulations (32, 33), results in no observable differences in pore radius and minimal area expansion (Fig. 1c). Increasing the tension beyond 40 mN/m appears to initiate expansion of the MscL pore, yet the very high tension destabilizes and ruptures the membrane preventing continuation of the MD simulation (Fig. 1c). We see that far-field tension, effectively pulling on the edges of the simulated membrane, produces not opening but a different ‘funneling’ transition when compared to the effect of tension distributed locally around the protein. We presume that the relatively soft lipid bilayer limits transmission of tension to the stiff protein, which possibly dampens productive fluctuations that would drive the outward motion of the helices.

As with any steered method, there are many ways to construct and bias the CV to reach the desired system configuration. The results shown in Fig. 1 where obtained using a linearly moving harmonic restraint or constant velocity pulling method where the biasing potential,

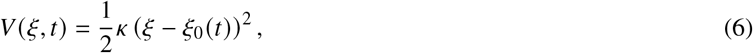

changes with time as the equilibrium position, *ξ*_0_(*t*), is gradually adjusted at a constant rate (*v*), *ξ*_0_(*t*) = *ξ*_0_(0) + *vt*. Constant velocity steering allows gentle gating of MscL as the lipid pulling forces start small and increase beyond the threshold needed to open the channel. Alternatively, one can apply a force of constant magnitude on the CV regardless of the value. We characterize the gating process through both of these strategies in the next section.

### MscL gating pathway

Having established an effective and systematic method to actuate MscL in MD, we now characterize the physical and structural properties of gating and compare to known experimental features from the literature. Rather than describe the actuation process in terms of the LDT CV, *ξ*, we use the change in projected protein area, Δ*A* = *A* – *A*_0_, as a more natural reaction coordinate that physically captures the current state of the channel at any point during gating, has a unique correspondence with *ξ* (Fig. 1b), and has been used as a parameter to analyze experimental patch-clamp data (25, 78). In the region of area expansion at Δ*A* = 20 nm^2^, as determined by patch-clamp predictions of the fully open state (25, 78), our simulation-predicted radius of 1.3 ± 0.1 nm is in good agreement with experiments that indicate a pore size of 2.5 – 3.0 nm (37, 49–52) and estimated conductance of 3.0 ± 0.1 nS (see Methods) agrees well with the experimental value of 3.2 nS (78) (see Fig. S2 in the Supporting Material).

The opening transition in MscL has been suggested to proceed through tilting of the TM helices resulting in expansion of the barrel accompanied by solvation of the hydrophobic constriction. Therefore, in the LDT-MD expansion trajectories we analyzed pore radius, tilting and crossing angles of TM helices, TM1-TM2 electrostatic energies as well as the number of water molecules in the vicinity of V21 (Fig. 2). We tested a variety of pulling velocities allowing complete transition to the open state in as fast as 4 ns to as slow as 100 ns. The force acting on the LDT CV during each simulation, 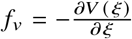, grows monotonically with area expansion (Fig. 2a) as expected, and it is generally larger for faster pulling rates given the non-equilibrium process where energy dissipation due to friction increases. MscL’s pore radius appears to correlate approximately linearly with area expansion except at the initial stage of gating (Δ*A* ≤ 5 nm^2^) where the radius changes much more slowly (Fig. 2b). This trend also appears to be unaffected by the timescale of the transition. More interestingly, the change in tilt angle of the TM helices with respect to the direction normal to the membrane (z) displays two distinct regimes as shown in Fig. 2c for TM1, 〈Δ*θ*_1,z_ (Δ*A*) = *θ*_1,z_ (Δ*A*) – *θ*_1,z_ (0)〉 (data averaged over all five subunits). During the initial channel expansion, Δ*A* = 0 – 10nm^2^, the TM helices undergo a large tilt of ≈ 20° that is accompanied by an increase in the crossing angle between TM1 and TM2 of the same subunit (Δ*θ*_1,2_, Fig. 2d). As TM1’ of the adjacent subunit further wedges in-between TM1 and TM2 during this initial transition, the TM1-TM2 short-range electrostatic interaction energy (*r_c_* ≤ 1.6 nm, Fig. 2e) arising largely from hydrogen-bonding interactions also significantly decreases.

**FIGURE 2.**
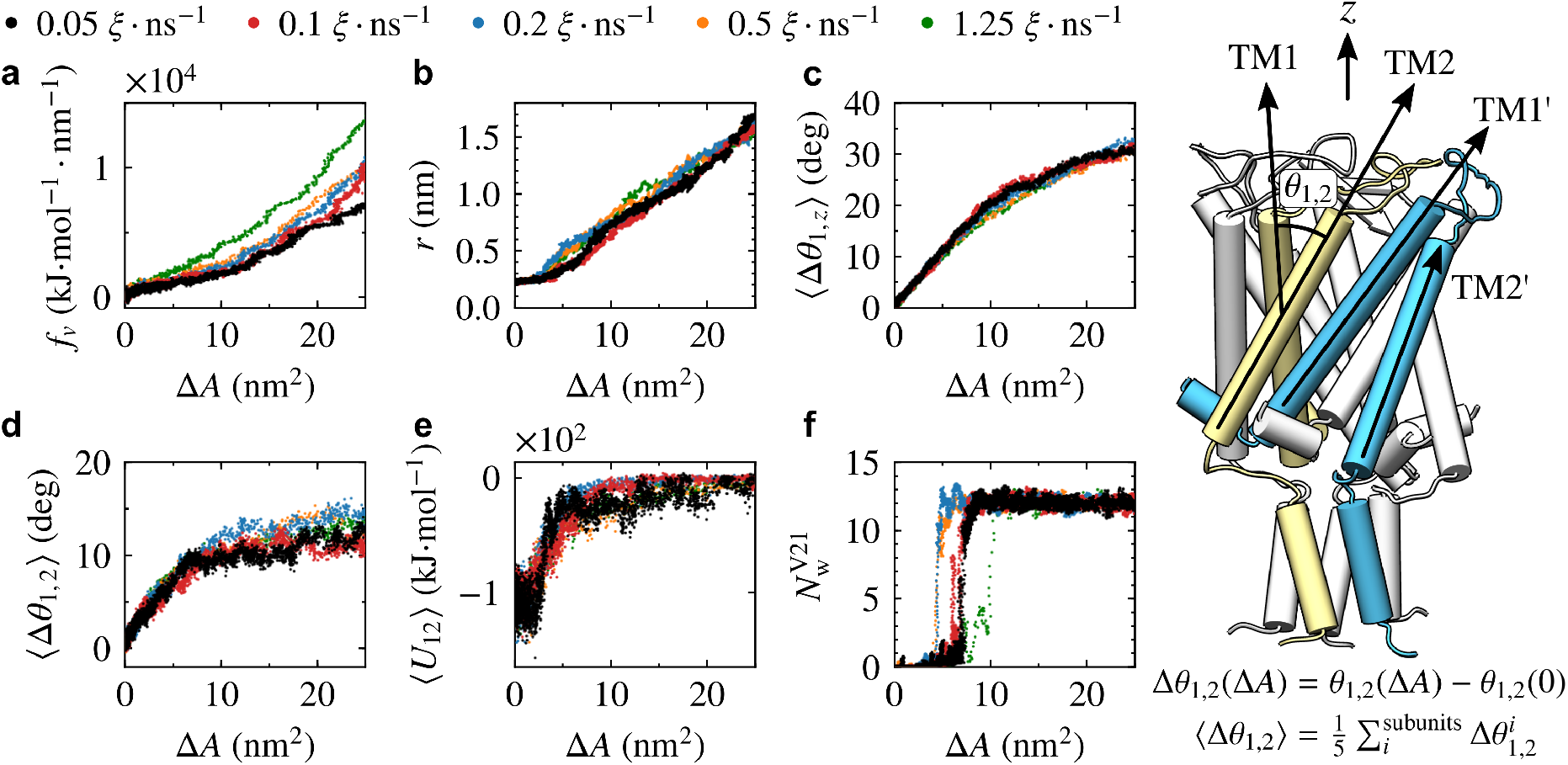
Structural properties of MscL during gating by constant velocity LDT-MD simulations as a function of protein area expansion (Δ*A*). Five velocities were tested: 0.05 (black), 0.1 (red), 0.2 (blue), 0.5 (orange), and 1.25 (green) *ξ* · ns^−1^. (a) Force acting on the LDT CV. (b) Pore radius, *r* (measured at residue V21). (c) Change in the tilt angle of TM1 relative to the direction normal to the plane (*z*). (d) Change in the crossing angle between TM1 and TM2 of the same subunit. (e) TM1-TM2 short-range electrostatic interaction energy (within a 1.6 nm cutoff). Quantities in panels (c), (d), and (e) are averaged values over the five channel subunits. (f) Number of water molecules within 0.4 nm of the center of mass of the five V21 residues. Illustration on the right shows schematically the TM helices and angles measured.

As the TM helices tilt and the pore expands, the ‘vapor lock’ formed by hydrophobic residues in the vicinity of V21 begins to wet and allows passage of water through the channel. In the constant-velocity LDT-MD simulations, wetting of the hydrophobic pore takes place at an area expansion of Δ*A* = 5 – 10 nm^2^ as measured by the number of waters within 0.4 nm from the center of mass of the five V21 residues (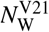, Fig. 2f). Further expansion of the channel between Δ*A* = 10 – 20 nm^2^ shows a more gradual tilt of the TM helices and much smaller change in the TM1-TM2 crossing angle. The large tilt of 30° in the vicinity of the open state (Δ*A* = 20 nm^2^) is in good agreement with results from electron paramagnetic spectroscopy experiments (50). The large tilting of the TM helices before wetting of the pore and subsequent outward motion is consistent with a silent pre-expansion of the closed state (79).

In comparison to constant velocity, LDT-MD simulations under constant pulling force show behavior analogous to the membrane tension simulations where at low LDT force there is minimal protein area expansion (Fig. 3a) and no visible change in the pore radius (Fig. 3b), while at larger forces the channel expansion occurs very quickly to a force-dependent maximum. As the pulling force increases, the protein area expansion where the pore radius steadily begins to grow is shifted to larger values of Δ*A* (Fig. 3c) and so is wetting of the hydrophobic pore as shown in 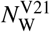 (Fig. 3d). Tilting of the TM helices is also shifted to larger values of Δ*A* with increasing pulling force (Fig. 3e) although the pattern in the TM1-TM2 crossing angle (Fig. 3f) appears to be largely unaffected and is similar to that observed in the constant velocity simulations. Together, these results indicate a kinetically-dependent effect where a rapidly occurring gating transition in MscL favors lateral expansion of the protein before the TM helices begin to noticeably tilt and the pore radius sufficiently expands to allow channel conduction.

**FIGURE 3.**
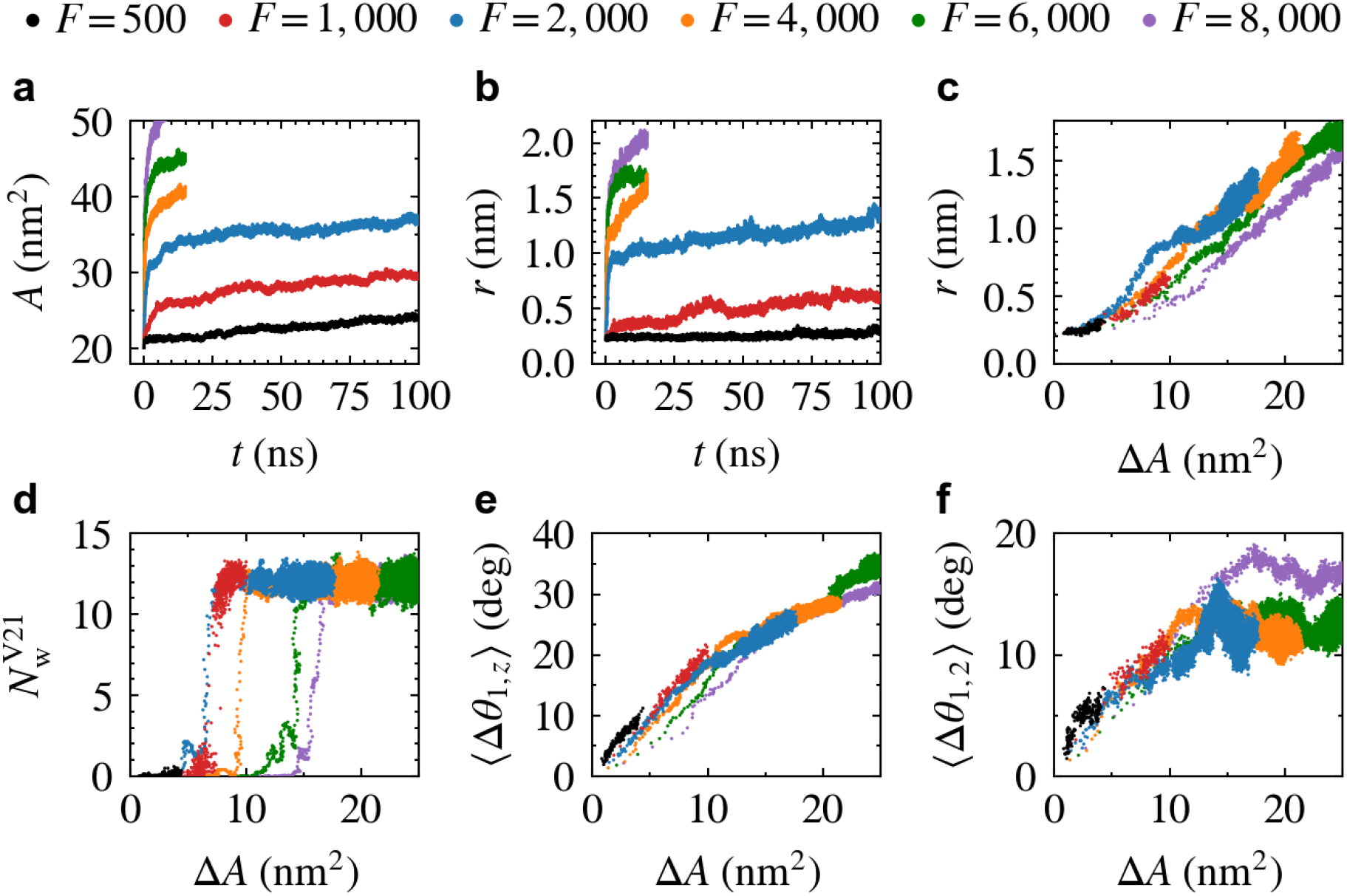
Structural properties of MscL during gating by constant force LDT-MD simulations. Six forces were tested: 500 (black), 1,000 (red), 2,000 (blue), 4,000 (orange), 6,000 (green), and 8,000 (purple) kJ · mol^−1^ · nm^−1^. (a) Protein in-plane projected area versus time. (b) Pore radius at residue V21 versus time. (c) Pore radius versus area expansion. (d) Number of water molecules within 0.4 nm of the center of mass of the five V21 residues. (e) Change in the tilt angle of TM1 relative to the direction normal to the plane (z). (f) Change in the crossing angle between TM1 and TM2 of the same subunit.

While the timescale of the gating process under conventional experimental conditions is likely to be much longer, in the order of microseconds (80), both the constant-velocity and constant-force simulations indicate that expansion of the pore, preceded by helical tilting, can be a gradual process guided not only by the tension acting on the membrane but also by intra-protein interactions that determine the motion of the TM helices. Pore wetting, however, is always a step-like and essentially instantaneous process (Fig. 2f and Fig. 3d). It was suggested previously (26) that synchronization of the main gating transition in WT MscL occurs through a single wetting event. Per-chain analysis shows uniform motion across all subunits during constant velocity gating in agreement with a synchronized transition (see Fig. S3 in the Supporting Material). Gain-of-function mutants with hydrophilic substitutions in the constriction are characterized by a permanently pre-wetted pore and exhibit a multitude of subconducting states that may be the result of asynchronous action of individual subunits (78). Systematic characterization of substates in MscL by LDT-MD is beyond the scope of this study and will be explored in future work.

### Energetics of opening under tension

Beyond structural characterization of MscL during opening, the LDT CV may be combined with enhanced sampling methods such as metadynamics (81, 82) or umbrella sampling (83) to estimate the energetics of gating. Membrane tension, *γ*, provides the energy needed for gating, in the form of Δ*E* = *γ*Δ*A*, and therefore we explore the free energy landscape under various tensions that stabilize the fully open state and potentially other substates. We combine the LDT CV with well-tempered metadynamics (65) (WTMetaD) to compute the free energy profiles for the MscL membrane system under three different membrane tensions of 10, 25, and 35 mN/m (below the 45 mN/m threshold where the membrane ruptures in our tests, see Fig. 1c) as well as under no tension. Although some of these values may seem excessively large given that the tension threshold for MscL is 10 – 15 mN/m (near the membrane lytic tension), finite size effects make it difficult to compare simulation and experimental tension values due to lack of long-range fluctuations in the orders of magnitude smaller simulation membrane patch (see Marsh for a detailed discussion (84)). At each tension, the membrane-protein system was first equilibrated followed by a constant velocity (*v* = 1.25 *ξ*/ns) LDT-MD simulation to generate the starting configurations of the multiple-walker WTMetaD calculation (200 walkers/tension) to obtain the estimated free energy as a function of the LDT CV, *ξ* (see Fig. S4 in the Supporting Material). Each walker was run for 30 ns for a combined simulation time of 6 *μ*s per tension.

As with the structural analysis in the previous section, we present the free energy profiles in Fig. 4a as a function of protein area expansion, Δ*A*, rather than the LDT CV for ease of comparison with experiments (see Fig. S4 for the correspondence between values of Δ*A* and *ξ*). The local energy minimum of the closed state at each tension is taken as the zero reference value for every curve in Fig. 4a. Convergence of the free energy calculations is shown in Fig. 4b by means of the root-mean-squared deviation as a function of the combined simulation time. In the absence of tension, the closed state sits in a deep energy well that keeps the channel in a very narrow range of areas (Fig. 4a, black line). A metastable state is observed at ~ Δ*A* = 4 nm^2^, likely due to rearrangement of the tightly packed TM helical bundle, but the energy continues to rapidly increase as the protein expands to larger areas. Application of net tension on the membrane produces various stable expanded MscL states in the range of Δ*A* = 5 – 20 nm^2^ (Fig. 4a). At the lowest tension tested, 10 mN/m, the energy rapidly grows to a maximum of 15 *k*_B_*T* for a modest increase in area (~ 2 nm^2^), similarly to the no tension curve. This maximum is then followed by a steady decrease in energy as the area continues to grow and multiple shallow local minima appear before the energy begins to quickly rise around Δ*A* > 12 nm^2^, at which point the membrane tension is unable to overcome the increasingly unfavorable energy of the expanding pore (Fig. 4a, red line).

**FIGURE 4.**
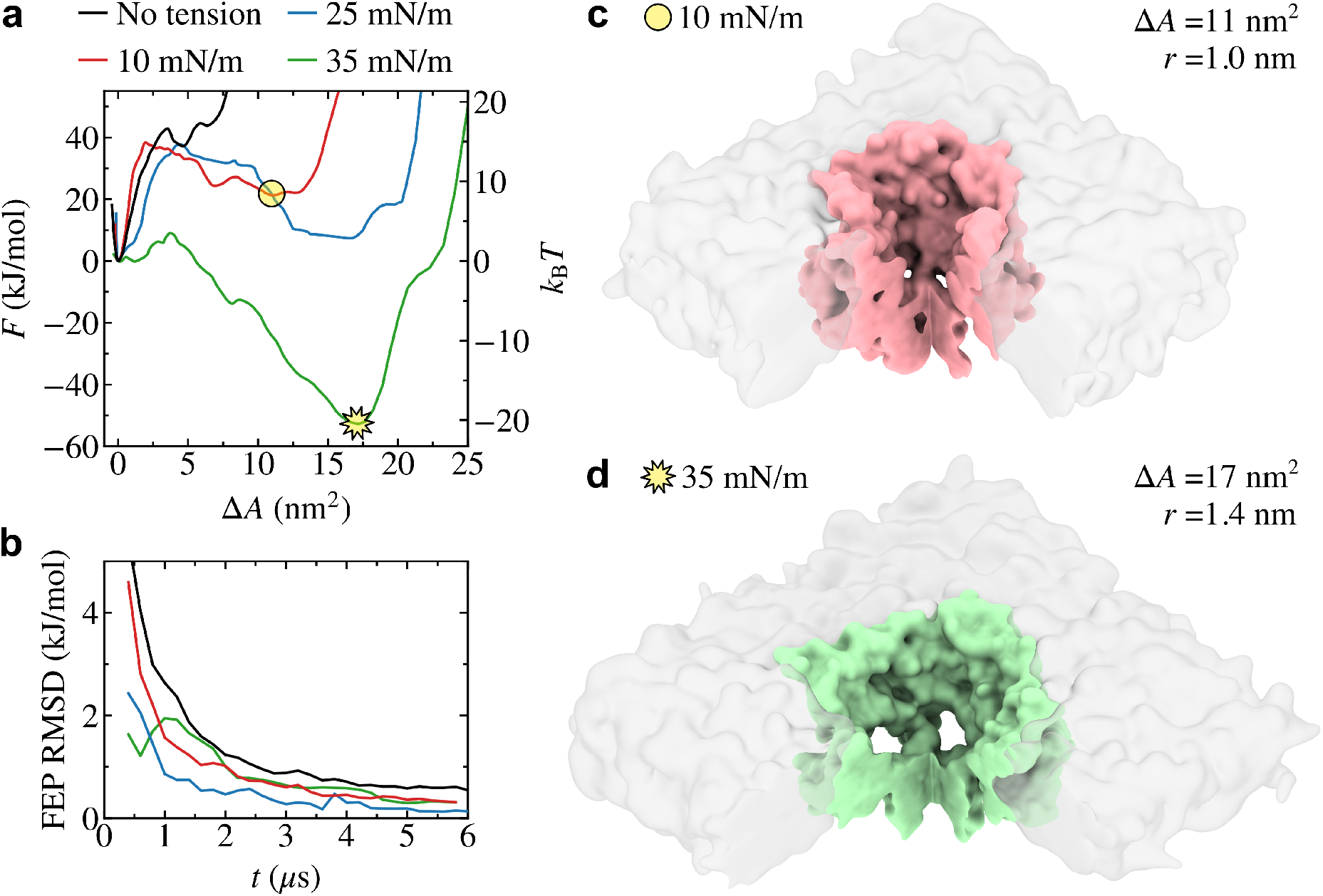
(a) Free energy profiles of MscL gating under tension obtained by combining the LDT CV with multiple-walker well-tempered metadynamics (see Methods). Although the free energy is computed as a function of *ξ* (see Fig. S4), we present it here as a function of the protein area expansion, Δ*A*, for ease of comparison with experiments. The local energy minimum of the closed state at each tension is taken as the zero reference value for every curve. (b) Convergence of computed free energy profiles measured by the root-mean-squared deviation as a function of the combined simulation time. Surface representations of expanded/open configurations near the local energy minimum at 10 mN/m (c) and 35 mN/m (d) shown on the right. Membrane colored in gray (transparent) and MscL colored in red or green with bottom quarter of the membrane/protein cut and removed to show pore opening.

The expanded configuration at Δ*A* = 11 nm^2^ observed for the 10 mN/m tension is not the fully open channel but likely corresponds to one of the experimentally observed substates as conduction would be largely hindered by the C-terminal bundle despite the relatively large pore radius, *r* = 1.0 nm (Fig. 4c). Increasing the tension to 25 mN/m stabilizes a fully open configuration in a shallow energy minima, of roughly equal energy as the closed state, at an area expansion of Δ*A* = 17 nm^2^ and with a pore radius of *r* = 1.4 nm (Fig. 4a, blue line). Further increasing the tension to 35 mN/m brings this same open state (Fig. 4d), with equal values of Δ*A* = 17 nm^2^ and *r* = 1.4 nm as observed for 25 mN/m, into a deep well that is 20 *k*_B_*T* lower in energy than the closed state (Fig. 4a, green line). The measured area expansion and pore radius of the common open state found in the 25 and 35 mN/m free energy profiles is in excellent agreement with experimental measurements of WT MscL (25, 37, 49–52, 78). The energy differences between states and corresponding barriers reported here are well within the range observed in the literature as conditions such as membrane thickness can significantly alter these values (25, 44, 78).

### Activation by asymmetric stimuli

As discussed in the Introduction, MscL can be gated by asymmetric addition of amphipathic molecules such as wedge-shaped lysolipids in the absence of membrane tension, i.e., incorporated to only one side of the membrane. We explore the role of asymmetric stimulus on the membrane by extending our LDT method to decouple contributions from the two membrane leaflets by separately grouping lipids in the periplasmic and cytoplasmic sides,

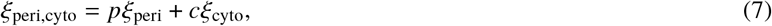

where the *p*:*c* ratio determines whether forces are larger on the periplasmic side (*p/c* > 1), cytoplasmic side (*p/c* < 1), or equally distributed (*p/c* = 1, as explored in the previous two sections). Constant velocity (*v* = 1.25 *ξ*/ns) LDT-MD simulations with varying *p:c* ratios show that gating can be induced by a wide range of stimuli with different levels of effectiveness (Fig. 5). Application of LDT with a larger cytoplasmic contribution, *ξ*_0.0,1.0_ and *ξ*_0.2,0.8_, shows similar increase in the pore radius versus protein area expansion compared to symmetric pulling, while LDT with a higher periplasmic contribution, *ξ*_0.8,0.2_, results in larger area expansion before the pore radius grows, and a purely periplasmic LDT, *ξ*_1.0,0.0_, is unable to open the pore (Fig. 5a). The force exerted on the LDT CV versus Δ*A* shows that the purely symmetric pull requires the lowest force to induce the large area expansion required for conduction, while forces for purely periplasmic or cytoplasmic LDT rapidly grow once the protein reaches a threshold around Δ*A* = 5 – 10 nm^2^ (Fig. 5b). Expansion profiles showing the subunit-averaged change in distance of alpha-carbons (Δ*r_cα_*) for all protein residues in the open state relative to the unbiased, equilibrium, configuration indicate that the shape of the open channel is strongly dependent on the ratio of periplasmic vs. cytoplasmic stimuli (Fig. 5c). Open configurations for each LDT-MD simulation in Fig. 5c were selected when the Δ*r_cα_* value for residue I23 was 1.2 nm to enable systematic comparison with single molecule FRET experiments of liposome-reconstituted EcMscL gating by lysolipids (51).

**FIGURE 5.**
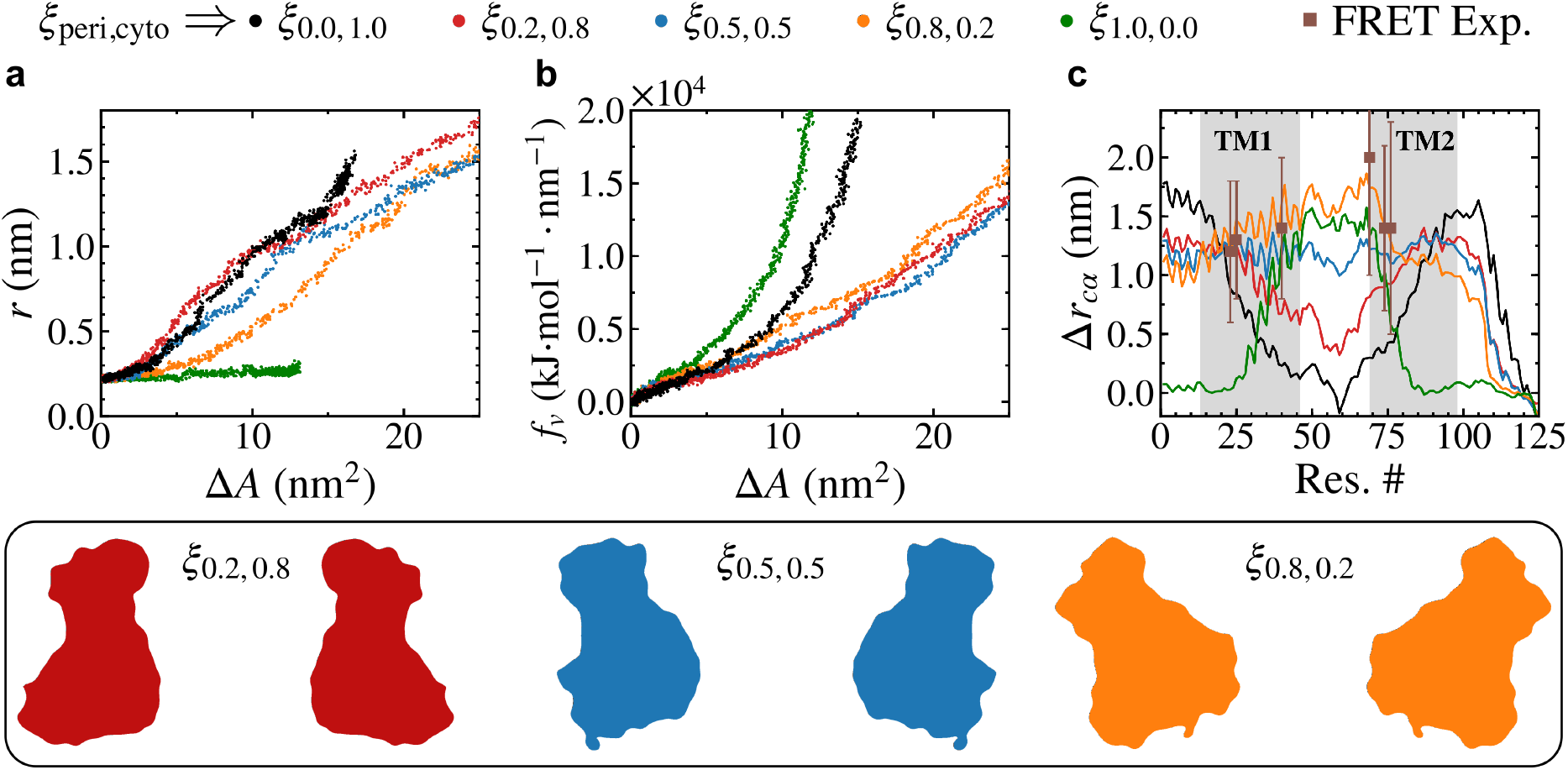
MscL activation by asymmetric constant velocity LDT-MD simulations with varying ratios of periplasmic to cytoplasmic LDT bias, *ξ*_peri,cyto_. Five stimuli with varying *p*:*c* ratios were tested: *ξ*_0.0,10_ (black), *ξ*_0.2,0.8_ (red), *ξ*_0.5,0.5_ (blue), *ξ*_0.8,0.2_ (orange), and *ξ*_1.0,0.0_ (green). Panels (a) and (b) show the pore radius (*r*) and biasing force (*f_v_*) versus protein area expansion (Δ*A*) respectively. (c) Expansion profiles showing the subunit-averaged change in distance of alpha-carbons (Δ *r_cα_*) for all protein residues in the open state (chosen when the Δ *r_cα_* value for residue I23 was 1.2 nm) relative to the unbiased configuration. Single molecule FRET experimental distances from Weng et al. (51) shown in brown squares. Bottom inset shows cross-sections of axis-symmetric density maps (including only the TM residues) obtained from the same protein configurations used for the analysis in panel (c).

Differences in the shape of the open channel under different stimuli are illustrated pictorially on the bottom inset of Fig. 5, where the colored silhouettes show cross-sections of axis-symmetric density maps (including only the TM residues) obtained from the same snapshots used for the analysis in Fig. 5c. The Δ*r_cα_* values for the periplasmic-focused LDT-MD stimuli, *ξ*_peri,cyto_ = *ξ*_0.8,0.2_ (Fig. 5c, orange line), show best agreement with the single molecule FRET experimental data from Weng et al. (51) (Fig. 5c, brown squares), where lysolipids were added only to the periplasmic side of the membrane. Comparison with our LDT-MD simulations suggests that one-sided incorporation of lysolipids generates an effective force that radially pulls on the channel on the same side of addition, and which may occur due to asymmetric membrane expansion. While one may naively conclude that increased crowding on the side of lysolipid addition would induce compression of the channel, the differential stress between the two leaflets, established by the asymmetric lipid distribution, may actually promote expansion of the crowded side of the membrane. This somewhat counter-intuitive phenomena has been demonstrated in coarse-grained MD simulations of asymmetric addition of lysolipids to planar bilayers (85).

## CONCLUSIONS

Through careful characterization and comparison with experimental observations, we have shown that the Locally Distributed Tension LDT-MD method is a powerful new tool to systematically and rapidly explore structure, dynamics, and energetics of the transduction process in mechanically activated membrane proteins such as MscL, while preserving the structure of the bilayer and protein-lipid contacts, which are crucial for a realistic force transduction and protein’s mechanical response. The excellent experimental agreement in the observed structural properties of MscL’s open state show that LDT-MD, in combination with a detailed atomistic simulation model, can directly provide accurate and physically meaningful results without needing to impose experimental constraints. Furthermore, it provides a flexible approach to investigate any MS membrane protein through symmetric and asymmetric stimuli by means of conventional steered MD or enhanced sampling free energy methods. Gating of MscL through LDT-MD simulations with various ratios of periplasmic/cytoplasmic bias provides new insights into the puzzling activation by addition of lysolipids, and suggests that a range of stimuli may actuate the channel albeit with different structural properties.

## SUPPORTING MATERIAL

Supporting Material can be found online at http://www.biophysj.org.

## AUTHOR CONTRIBUTIONS

RRT and JMV setup and carried out MD simulations. RRT and JMV performed data analysis and created figures. AA provided conductance analysis scripts. JMV conceived the project and designed simulation studies with input from RRT, AA, and SS. RRT, AA, SS, and JMV contributed to writing of paper, reviewed, and approved it in its final form.

## ACKNOWLEDGMENTS

JMV acknowledges the support of the National Science Foundation through Grant No. CHE-1944892. Computations were performed, in part, on the Vermont Advanced Computing Core supported in part by NSF Award No. OAC-1827314.

